# First record of the invasive red lionfish (*Pterois volitans*) in the Parcel de Manuel Luís Marine State Park, Brazilian Equatorial Margin

**DOI:** 10.1101/2025.07.25.666735

**Authors:** Marcelo Henrique Lopes Silva, James Werllen de Jesus Azevedo, Leonardo Silva Soares, João Luiz Baptista de Carvalho, Filipe França dos Santos Silva, Francisco das Chagas Miranda de Carvalho Neto, Nicollas Silva Mendes, Antonio Carlos Leal de Castro

## Abstract

The red lionfish (*Pterois volitans*), native to the Indo-Pacific, is recognized as one of the most harmful invasive species in the marine ecosystems of the Western Atlantic. Since its accidental introduction, the species has rapidly expanded across the Caribbean, Gulf of Mexico, and Brazilian coast, reaching marine protected areas with high ecological value. This study documents the first confirmed occurrence of *P. volitans* in the reef environment of the Parcel de Manuel Luís Marine State Park, located on the Brazilian Equatorial Margin. The record was based on underwater footage collected during a scientific expedition in June 2025. Two individuals were observed near the wreck of the ship Ana Cristina at a depth of five meters. The specimens exhibited typical morphological features and behavioral patterns associated with foraging and habitat exploration. Physicochemical variables measured at the site confirmed stable tropical marine conditions, including high salinity and temperature, compatible with the ecological tolerance of the species. Additional measurements at eight surrounding stations revealed minimal stratification, reinforcing the environmental suitability for the species. Statistical analysis detected no significant differences in salinity or dissolved oxygen between surface and bottom waters, and a slight variation in temperature. The presence of *P. volitans* in a conservation unit raises concern about its potential expansion into other reef and coastal environments. Given its high reproductive capacity, absence of natural predators, and ecological plasticity, the species poses a serious threat to native biodiversity and trophic dynamics. The results highlight the urgency of establishing systematic monitoring programs and early-response strategies for invasive species in Brazilian reef systems.

## INTRODUCTION

The red lionfish (Pterois volitans), which is native to the Indo-Pacific, is recognized as one of the most impactful invasive species in marine ecosystems of the Western Atlantic Ocean (Hwang et al., 2025; del Rio et al., 2023). Since its accidental introduction in the 1980s and 90s, the species has spread rapidly through oceanic larval dispersal and anthropogenic introduction, colonizing large areas of the Caribbean Sea, Gulf of America, and, more recently, the Brazilian coast (Schofield, 2009; Morris & Whitfield, 2009; Rocha et al., 2015; Ferreira et al., 2019). This expansion is favored by the characteristics of the species, such as high fecundity, a generalist diet, the absence of effective natural predators, and broad physiological tolerance (Albins & Hixon, 2008; Côté & Smith, 2018; Green et al., 2012).

The first confirmed records in Brazil occurred in 2014 in the Fernando de Noronha Archipelago (Ferreira et al., 2015), followed by reefs and artificial structures along the coast of the North and Northeast regions of the country, indicating a pattern of progressive advance towards the Brazilian Equatorial Margin (Sampaio et al., 2022; Pinheiro et al., 2023; Ferreira et al., 2025). Recent ecological modeling suggests that the species will continue to expand its range along the Brazilian coast, including marine conservation units (Soares et al., 2025). The authors cited stress the vulnerability of protected areas among the states of the North and Northeast regions and project that, without effective control measures, the species will occupy the entire coastline by 2030. Maranhão was one of the states with no official records.

The invasion of *P. volitans* has caused serious ecological impacts in various locations, with significant declines in the abundance of native species, especially small fishes and juveniles, altering the trophic structure and reducing the resilience of reefs (Albins, 2013; Green & Côté, 2009; Lesser & Slattery, 2011). The presence of the species in protected areas demonstrates its high adaptability and underscores the need for specific management and monitoring measures, even in areas with legal conservation status.

On the coast of the state of Maranhão, the Parcel de Manuel Luís (PML) Marine State Park, which is approximately 180 km offshore, houses one of the largest reef formations of the Brazilian Equatorial Margin (Amaral et al., 2006). Despite its ecological importance and protected status, surveillance efforts targeting invasive species are incipient, increasing the vulnerability of the system to initial colonization events.

The present study documents the first confirmed record of *Pterois volitans* on the coast of Maranhão based on underwater images obtained within the PML Marine State Park. This finding expands the known geographic distribution of the invasive species in the Western South Atlantic and underscores the urgent need to implement continuous monitoring, prevention, and containment programs for the species in vulnerable reef ecosystems.

## MATERIAL AND METHODS

### Study area

The study was conducted from June 16 to 20, 2025, at PML Marine State Park, which is a protected marine conservation unit located on the continental shelf of the Brazilian Equatorial Margin, approximately 180 km off the coast of the state of Maranhão in northeastern Brazil (Figure 1). The protected area covers approximately 45,937 hectares and is composed of an extensive submerged reef formation, with depths ranging from five to 30 meters. The predominant substrate includes coral reefs of calcareous algae and hermatypic corals, associated with sandy bottoms and consolidated bioconstructed structures. This structural heterogeneity favors high biodiversity, especially of reef fishes and pelagic species.

**Fig. 1:**
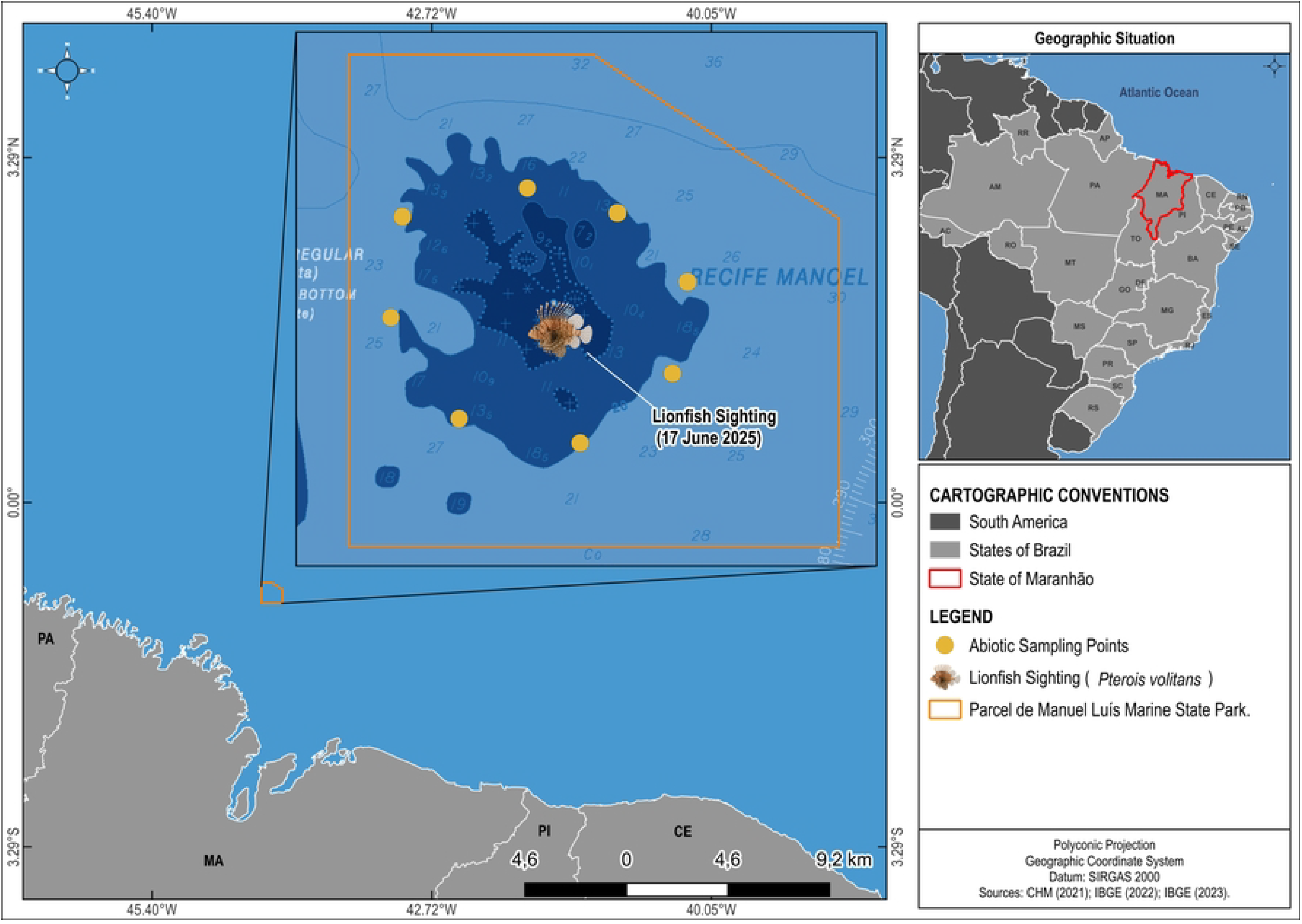
**Location of Parcel de Manuel Luís Marine State Park, highlighting the point of the record of *Pterois volitans* and stations for the collection of abiotic data obtained with a CTD device during the expedition in June 2025.**

Observations were performed in the vicinity of the wreck of the ship named Ana Cristina (00°52’43” S; 44°15’50” W), which occurred in 1984. The site is characterized by artificial structures covered with benthic organisms and constitutes a recurring sampling point in scientific studies due to its accessibility and the high diversity of species.

### Procedures and analyses

The record of the species *Pterois volitans* was made during scuba diving using the rapid visual technique, following the protocol described by Kimmel (1985). Filming took place during a 10-minute travel in the vicinity of the wreck, using a GoPro Hero 11 Black camera, which was manually operated by an experienced diver. The equipment was configured for 4K resolution to optimize image quality.

The underwater images were subsequently analyzed using EventMeasure software (v3.51; SeaGIS Pty. Ltd.), enabling the recording of the precise time of observation and the proportional measurement of the individuals. Taxonomic identification on the species level was based on diagnostic morphological characteristics (Schofield, 2009; Ferreira et al., 2019), such as color pattern, number and arrangement of body stripes, fin morphology, and head proportion. Identification was confirmed by experts experienced in visual recordings of *P. volitans*.

Environmental data were obtained using a conductivity-temperature-depth (CTD) profiler to characterize the oceanographic conditions of the area at the time of the sighting. Measurements were taken during the same dive at a depth of approximately five meters and included the following variables: water temperature (°C), salinity, dissolved oxygen (mg/L), turbidity (NTU), and depth (m).

Additional sampling was conducted using a CTD Rosette at eight different points around the PML Marine State Park. Physicochemical water variables were measured at two depths: surface (≈3 meters) and bottom (14 to 46 meters). The variables were water temperature (°C), salinity, dissolved oxygen (mg/L), pH, electrical conductivity, and redox potential (Eh, mV). The data were processed in a laboratory environment, following the operating protocols of the equipment used.

Mean and standard deviation values were calculated for each variable to describe variability among the sampling points. The Mann-Whitney test was used to test differences in values measured at the surface and bottom, adopting a 5% significance level (α = 0.05). Statistical analyses were performed using the R software (version 4.3.2).

The presence of the species was confirmed based on the MaxN criterion (Cappo et al., 2004), corresponding to the largest number of specimens observed simultaneously in a single frame of the footage – in this case, a single specimen. Total length was estimated visually by proportional comparison to elements of the environment and based on the camera scale, considering 5-cm increments. Additional information on the morphology and ecology of the species was obtained from Pinheiro et al. (2018) and Froese & Pauly (2025).

The activities were carried out within the scope of the project entitled “Oceanographic Bases for the Conservation of the Parcel Manuel Luís State Marine Park”, with funding from the State of Maranhão Research and Scientific and Technological Development Foundation (FAPEMA) and with the authorization of the State Secretary of the Environment and Natural Resources (SEMA/MA).

## RESULTS

During a scientific expedition conducted in June 2025 in the PML Marine State Park, two individuals of the species *Pterois volitans* were recorded associated with artificial reef structures formed by the wreck of the ship Ana Cristina (Figure 2). The specimens exhibited the typical color pattern of the species, with alternating vertical red and white stripes, an elongated body, and expanded dorsal and pectoral fins with prominent spines.

**Fig. 2:**
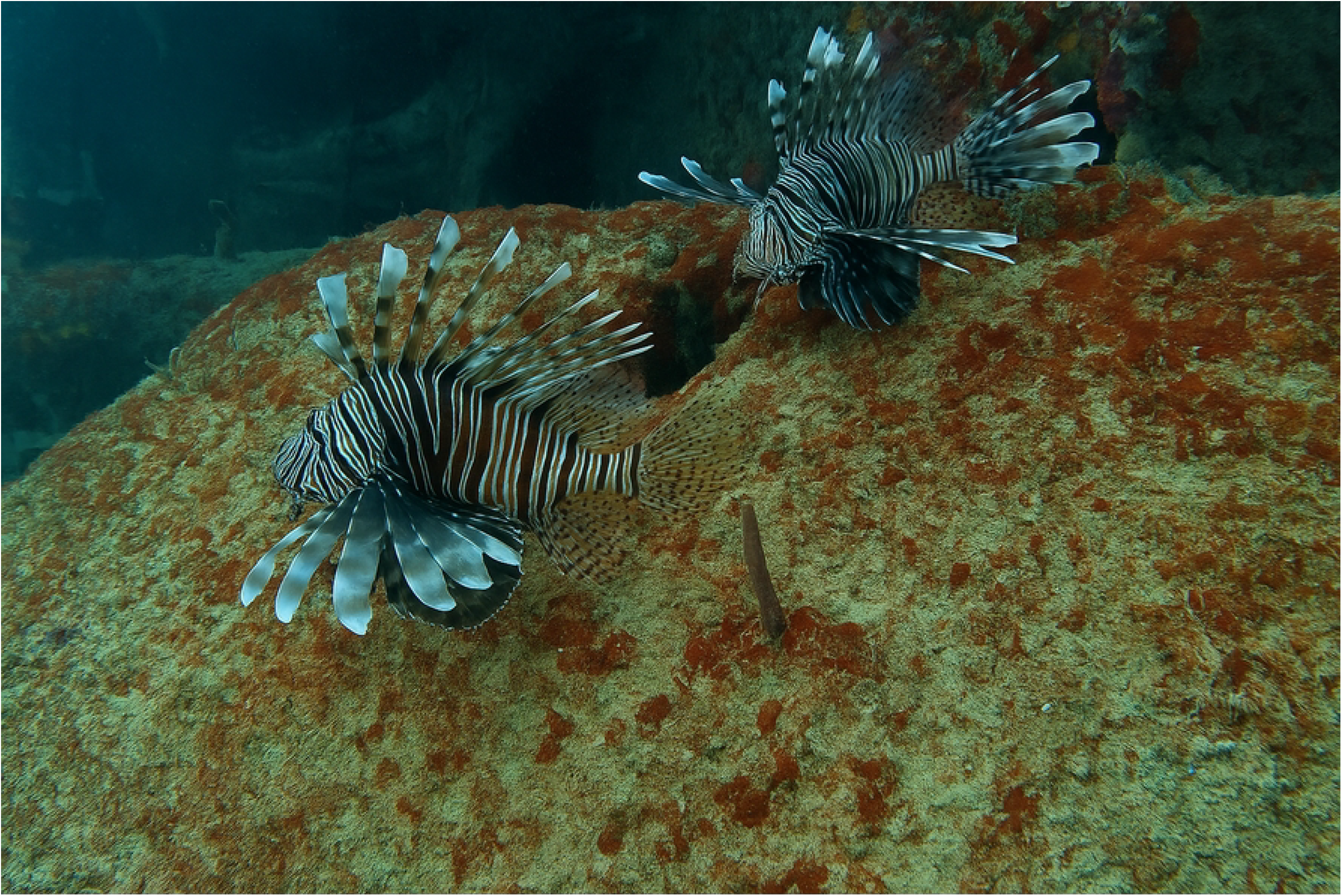
**Specimens of *Pterois volitans* recorded in the reef area of Parcel de Manuel Luís Marine State Park at a depth of 5 m during a scientific expedition from June 16 to 20, 2025.**

Visual estimates of total length were between 15 and 20 cm, suggesting subadults or young adults. Both individuals were observed actively moving across the surface of the wreck, exploring cavities, recesses, and areas covered in macroalgae and coral. During filming, the specimens were in an upright posture with their fins spread open and did not demonstrate evasive behavior due to the approach of the diver, remaining visible throughout the entire observation. The behavior resembled patterns of patrolling and the exploration of microhabitats, which are characteristics of foraging activities. Such observations suggest the use of the environment as a shelter and possible feeding grounds.

The physicochemical conditions recorded during the dive in the wreck area revealed a typically oligotrophic, thermally stable environment. At an approximate depth of five meters, water temperature was 28.37°C, salinity was 36.34, dissolved oxygen was approximately 5.6 mg/L, and density was 1,023.3 kg/m^³^. The absolute pressure was approximately 5.6 dbar. These values indicate a stable marine environment, with high salinity and temperature compatible with tropical reef zones, providing favorable conditions for the occupation of exotic species, such as *Pterois volitans*.

The physicochemical variables of the water were measured at the surface (≈3 meters) and bottom (14 to 46 meters in depth) at eight points distributed around the PML Marine State Park. The mean and standard deviation values of dissolved oxygen (DO), salinity, and temperature are displayed in Table 1. The values exhibited little variation between the strata of the water column. Additionally, pH, conductivity, and water density (σt) were analyzed and included in the statistical assessment.

**Table 1.** Mean and standard deviation values of physicochemical variables at surface (∼3 m) and bottom (14 to 46 m) at eight points around Parcel de Manuel Luís Marine State Park in June 2025.

| Variable | Surface (n = 8) | Bottom (n = 8) |
| --- | --- | --- |
| Dissolved oxygen (mg/L) | 3.54 ± 0.36 | 3.46 ± 0.25 |
| Salinity | 34.86 ± 1.34 | 34.70 ± 1.20 |
| Temperature (°C) | 23.05 ± 2.19 | 23.00 ± 2.07 |

The mean dissolved oxygen concentration was slightly higher at the surface (3.54 ± 0.36 mg/L) compared to the bottom (3.46 ± 0.25 mg/L). Salinity was similar between the two layers (34.86 ± 1.34 at the surface and 34.70 ± 1.20 at the bottom). Water temperature also varied little between the strata (23.05 ± 2.19 °C at the surface and 23.00 ± 2.07 °C at the bottom). The results of the Mann-Withney test revealed no statistically significant differences in dissolved oxygen (p = 0.798), salinity (p = 0.372) or (p = 0.344) between the surface and bottom, indicating a vertically homogeneous water column at the time of sampling.

## DISCUSSION

The record of *Pterois volitans* within PML Marine State Park constitutes the first confirmed report of the invasive species in the Brazilian Equatorial Margin, expanding its distribution beyond previously documented areas on the northern coast of Brazil. This finding anticipates recent projections, such as those by Soares et al. (2025) and Ferreira et al. (2025), who predicted the colonization of marine conservation units on the coast of the North and Northeast regions of the country by 2030 based on ecological modeling and larval dispersal.

The presence of this species in a reef conservation unit with considerable structural complexity is consistent with records from other tropical regions (Côté et al., 2013; Green et al., 2012), in which natural and artificial reefs are preferentially occupied due to the abundance of places to find shelter and prey items. In the PML Marine State Park, the individuals actively explored the wreck of the ship Ana Cristina, which is consistent with behavioral patterns described by Luiz et al. (2025) and Côté & Maljkovic (2010), who reported the association of the species with prominences and cavities used as foraging and shelter zones.

From the ecological standpoint, *P. volitans* is recognized as an opportunistic species with broad behavioral and physiological plasticity. Its ability to tolerate environmental variations is well documented (Kimball et al., 2004; Kulbicki et al., 2012) and the physicochemical data collected during the expedition reinforce this characteristic. At the location of the sighting, salinity was 36.3, temperature ranged from 28.4 to 28.6°C, and dissolved oxygen was approximately 4.0 mg/L, which are consistent with values reported in invaded habitats in other tropical regions (Johnston & Purkis, 2015). These parameters do not constitute physiological barriers and favor the survival and eventual establishment of the species in the reef environment.

The statistical analysis of data collected at eight sites around the PML Marine State Park indicated no significant differences between surface and bottom in terms of salinity, dissolved oxygen and temperature. The relative hydrochemical stability may contribute to the success of invasive species, such as the red lionfish, by reducing environmental limitations and expanding the range of exploitable habitats (Biggs & Olden, 2011).

Although only two individuals were recorded in the present study, experiences in other regions indicate that even small initial populations can proliferate rapidly due to the high fecundity of the species as well as the lack of natural predators and effective predatory behavior (Albins & Hixon, 2008; Morris & Whitfield, 2009). The presence of *P. volitans* poses additional risks in the PML Marine State Park, which is home to vulnerable reef species of both ecological and economical importance, such as *Epinephelus itajara, Mycteroperca bonaci*, and *Scarus trispinosus*. The potential for niche overlap and competition for resources can trigger trophic imbalances and a reduction in the abundance of native species (Ingeman, 2016; Hackerott et al., 2013).

Given the location of the PML Marine State Park on the inner continental shelf and its connectivity with adjacent coastal systems, the presence of *Pterois volitans* in the region raises concerns with regards to its potential for dispersal into sensitive coastal environments. Studies conducted in other regions of the Western Atlantic Ocean demonstrate that the lionfish is capable of colonizing a wide variety of habitats, including estuaries, mangroves, and sheltered areas close to the mainland (Barbour et al., 2010; Jud et al., 2011; Biggs & Olden, 2011).

The coastal and marine environments of the northern coast of Brazil are known for high productivity and vulnerability to ecological disturbances. The arrival of *P. volitans* in these areas could compromise local biodiversity, affect native fish populations, and alter the trophic structure of these ecosystems. It is therefore essential to implement integrated monitoring and management strategies that are not limited to marine conservation units but encompass the entire coastal zone potentially susceptible to invasion.

In summary, the present record confirms the presence of *Pterois volitans* on a coastal reef strategically located in the Brazilian Equatorial Margin, expanding the known distribution of the species in the Western South Atlantic. Its occurrence in a marine conservation unit lends strength to projections of its expansion and underscores the risk of the colonization of protected areas by invasive species, demonstrating the need for intensified monitoring and the implementation of management strategies directed at the conservation of the biodiversity of Brazilian reefs.

## CONCLUSION

The first confirmed record of *Pterois volitans* on the coast of the state of Maranhão, specifically in the reef area of the Parcel de Manuel Luís Marine State Park, expands the known distribution of this invasive species in the Western South Atlantic and constitutes a warning sign for the region of the Brazilian Equatorial Margin. The occurrence of the species in a protected reef environment under abiotic conditions compatible with its broad ecological tolerance underscores its potential for establishment and dispersal into adjacent habitats, including coastal and estuarine ecosystems of considerable socio-environmental importance.

Given the threat that the red lionfish poses to marine biodiversity and the stability of reef systems, it is essential to implement continuous monitoring programs and coordinated management actions, with the participation of research institutions, local communities, and environmental agencies. Early detection of new sites of occurrence can assist in the establishment of public control policies to mitigate the ecological impacts documented in other invaded regions. The results presented here contribute to the understanding of the invasion dynamics of this species in Brazil and underscore the urgent need to expand environmental surveillance efforts throughout the northern and northeastern coasts of the country.

## ACKNOWLEDGMENTS

We thank the Foundation for Research and Scientific and Technological Development of Maranhão (FAPEMA) for funding this research. We also acknowledge the Maranhão State Secretariat for the Environment and Natural Resources (SEMA-MA) for issuing the research license. Special thanks to **Lieutenant José Barbosa Fonseca Neto**, stationed at the Search and Rescue Battalion (BBS) of the Maranhão State Fire Department, for his support in obtaining underwater footage during diving operations. The Coordination for the Improvement of Higher Education Personnel (CAPES) is also acknowledged for supporting the academic training of the authors.

## REFERENCES

Albins, M. A., & Hixon, M. A. (2008). Invasive Indo-Pacific lionfish Pterois volitans reduce recruitment of Atlantic coral-reef fishes. Marine Ecology Progress Series, 367, 233– 238.

Albins, M. A., & Hixon, M. A. (2013). Worst case scenario: potential long-term effects of invasive predatory lionfish (Pterois volitans) on Atlantic and Caribbean coral-reef communities. Environmental Biology of Fishes, 96, 1151–1157.

Barbour, A. B., Montgomery, M. L., Adamson, A. A., Díaz-Ferguson, E., & Silliman, B. R. (2010). Mangrove use by the invasive lionfish Pterois volitans. Marine Ecology Progress Series, 401, 291–294.

Biggs, C. R., & Olden, J. D. (2011). Multi-scale habitat occupancy of invasive lionfish (Pterois volitans) in coral reef environments. Environmental Biology of Fishes, 93, 451– 460.

Cappo, M., Speare, P., & De’ath, G. (2004). Comparison of baited remote underwater video stations (BRUVS) and prawn (shrimp) trawls for assessments of fish biodiversity in interreefal areas of the Great Barrier Reef Marine Park. Journal of Experimental Marine Biology and Ecology, 302(2), 123–152.

Claydon, J. A. B., Calosso, M. C., & Traiger, S. B. (2012). Progression of invasive lionfish in seagrass, mangrove and reef habitats. Marine Ecology Progress Series, 448, 119–129.

Côté, I. M., & Maljkovic, A. (2010). Predation rates of Indo-Pacific lionfish on Bahamian coral reefs. Marine Ecology Progress Series, 404, 219–225.

Côté, I. M., & Smith, N. S. (2018). The lionfish Pterois sp. invasion: Has the worst-case scenario come to pass?. Journal of Fish Biology, 92(3), 660–689.

Côté, I. M., Green, S. J., & Hixon, M. A. (2013). Predatory fish invaders: insights from Indo-Pacific lionfish in the western Atlantic and Caribbean. Biological Conservation, 164, 50–61.

Cure, K., McIlwain, J. L., & Hixon, M. A. (2014). Habitat plasticity in native and invasive coral-reef fishes. Marine Ecology Progress Series, 506, 219–229.

Ferreira, C. E. L., Luiz, O. J., Floeter, S. R., Lucena, M. B., Barbosa, M. C., Rocha, C. R., & Gasparini, J. L. (2019). First invasion of Indo-Pacific lionfish (Pterois volitans) in the South Atlantic Ocean. Biological Invasions, 17, 1387–1390.

Ferreira, C. E. L., Soares, M. O., & Luiz, O. J. (2025). Emerging invasion of lionfish (Pterois volitans) in Brazilian marine protected areas: projected range and ecological concerns. Marine Environmental Research, 200, 105544.

Frazer, T. K., Jacoby, C. A., Edwards, M. A., Barry, S. C., & Manfrino, C. M. (2012). Coping with the lionfish invasion: can targeted removals yield beneficial effects?. Reviews in Fisheries Science, 20(4), 185–191.

Green, S. J., & Côté, I. M. (2009). Record densities of Indo-Pacific lionfish on Bahamian coral reefs. Coral Reefs, 28(1), 107.

Green, S. J., Akins, J. L., Maljković, A., & Côté, I. M. (2012). Invasive lionfish drive Atlantic coral reef fish declines. PLoS ONE, 7(3), e32596.

Hackerott, S., Valdivia, A., Green, S. J., Côté, I. M., Cox, C. E., Akins, L., … & Bruno, J. F. (2013). Native predators do not influence invasion success of Pacific lionfish on Caribbean reefs. PLoS ONE, 8(7), e68259.

Hare, J. A., & Whitfield, P. E. (2003). An integrated assessment of the introduction of lionfish (Pterois volitans/miles complex) to the western Atlantic Ocean. NOAA Technical Memorandum NOS NCCOS, 2, 21.

Ingeman, K. E. (2016). Lionfish cause increased mortality in resident coral-reef fishes. Marine Ecology Progress Series, 558, 235–245.

Johnston, M. W., & Purkis, S. J. (2015). A coordinated and sustained international strategy is needed to turn the tide on the Atlantic lionfish invasion. Marine Ecology Progress Series, 533, 219–235.

Jud, Z. R., Layman, C. A., & Lee, J. A. (2011). Broad salinity tolerance in the invasive lionfish Pterois spp. may facilitate estuarine colonization. Environmental Biology of Fishes, 92(4), 511–516.

Kimball, M. E., Miller, J. M., Whitfield, P. E., & Hare, J. A. (2004). Thermal tolerance and potential distribution of invasive lionfish (Pterois volitans/miles complex) on the East Coast of the United States. Marine Ecology Progress Series, 283, 269–278.

Kimmel, J. J. (1985). A new species-time method for visual assessment of fishes and other mobile epibenthos in shallow habitats. U.S. National Park Service Technical Report.

Kletou, D., Hall-Spencer, J. M., & Kleitou, P. (2016). A lionfish (Pterois miles) invasion has begun in the Mediterranean Sea. Marine Biodiversity Records, 9(1), 46.

Kulbicki, M., Beets, J., Chabanet, P., Cure, K., Darling, E., Floeter, S. R., … & Galzin, R. (2012). Distributions of Indo-Pacific lionfishes (Pterois spp.) in their native ranges: implications for the Atlantic invasion. Marine Ecology Progress Series, 446, 189–205.

Luiz, O. J., Ferreira, C. E. L., & Floeter, S. R. (2025). Habitat selection and predatory behavior of invasive lionfish (Pterois volitans) in Brazilian reefs. Journal of Experimental Marine Biology and Ecology, 560, 111741.

Morris, J. A., & Whitfield, P. E. (2009). Biology, ecology, control and management of the invasive Indo-Pacific lionfish: an updated integrated assessment. NOAA Technical Memorandum NOS NCCOS 99.

Pinheiro, H. T., Ferreira, C. E. L., Joyeux, J. C., Santos, R. G., & Horta, P. A. (2018). Reef fishes of the Brazilian Exclusive Economic Zone: recent advances and baseline for conservation. Latin American Journal of Aquatic Research, 46(5), 845–869.

Soares, M. O., Ferreira, C. E. L., & Luiz, O. J. (2025). Potential spread of the invasive lionfish in Brazilian marine protected areas. Marine Ecology Progress Series, 722, 125–13.

